# Label-free quantitative proteomics analyses of mouse astrocytes provides insight into the host response mechanism at different developmental stages of *Toxoplasma gondii*

**DOI:** 10.1101/2023.01.15.524169

**Authors:** Huanhuan Xie, Chao Xu, Guihua Zhao, Hongjie Dong, Lisha Dai, Haozhi Xu, Lixin Zhang, Hang Sun, Qi Wang, Junmei Zhang, Kun Yin

## Abstract

*Toxoplasma gondii* (*T. gondii*) is an opportunistic parasite that can infect the central nervous system (CNS), causing severe toxoplasmosis and behavioral cognitive impairment. Mortality is high in immunocompromised individuals with toxoplasmosis, most commonly due to reactivation of infection in the CNS. There are still no effective vaccines and drugs for the prevention and treatment of toxoplasmosis. There are five developmental stages for *T. gondii* to complete life cycle, of which the tachyzoite and bradyzoite stages are the key to the acute and chronic infection. In this study, to better understanding of how *T. gondii* interacts with the host CNS at different stages of infection, we constructed acute and chronic infection models of *T. gondii* in astrocytes, and used label-free proteomics to detect the proteome changes before and after infection, respectively. A total of 4676 proteins were identified, among which 163 differentially expressed proteins (DEPs) (fold change≥1.5 or ≤0.67 and *p*-value≤0.05) including 109 up-regulated proteins and 54 down-regulated proteins in C8-TA vs C8 group, and 719 DEPs including 495 up-regulated proteins and 224 down-regulated proteins in C8-BR vs C8-TA group. After *T. gondii* tachyzoites infected astrocytes, DEPs were enriched in immune-related biological processes to promote the formation of bradyzoites and maintain the balance of *T. gondii*, CNS and brain. After *T. gondii* bradyzoites infected astrocytes, the DEPs up-regulated the host’s glucose metabolism, and some up-regulated DEPs were closely related to neurodegenerative diseases. These findings not only provide new insights into the psychiatric pathogenesis of *T. gondii*, but also provide potential targets for the treatment of acute and chronic Toxoplasmosis.

## 1. Introduction

*Toxoplasma gondii* (*T. gondii*) is an obligate intracellular parasite that can infect almost all warm-blooded animals, including humans and livestock, and causing zoonotic parasitic diseases^[1-4]^. It is estimated that 30% of the world population was infected by *T. gondii*, and in China, approximately 8.2% Chinese population was infected with it^[5,6]^. *T. gondii* has a complex life cycle, and almost all warm-blooded animals can become intermediate hosts, except for the terminal host felid. During the intracellular life, *T. gondii* undergoes five developmental stages including tachyzoite, bradyzoite, sporozoite, schizont, and gametocyte. Tachyzoites are parasitic in pseudocyst, and bradyzoites are present in the tissue cyst. The schizont and gametophyte are the sexual reproductive stages, and finally formed into oocyst. In the intermediate hosts, there are two infectious stages: tachyzoite and bradyzoite. Since the morphological difference between tachyzoite and bradyzoite is quite small, the bradyzoite-specific expression protein BAG1 is usually used for the identification of bradyzoites.

The mutual transformation of tachyzoites and bradyzoites is the central link in the pathogenesis of *T. gondii*. Tachyzoites can cause acute infection, while bradyzoites are the main cause of chronic infection. Acute toxoplasmosis is more prevalent in immunodeficiency patients^[7-9]^. However, when the tachyzoites are transformed into bradyzoites and form cysts in normal immune responses, they will latently parasitize in the host’s brain and ocular chorioretinal areas. When the hosts immune system is impaired, the latent bradyzoites (in the CNS) could burst out of the cysts, reconvert to replicative tachyzoites and triggering a new round of infection^[10]^. In the chronic stages of infection, the host may have alterations in behavior and cognition^[11-14]^. For example, in rodent hosts, the chronic infection can lead to excessive exercise, reduced anxiety, reduced new phobias, and predator vigilance. In particular, *T. gondii* could induce changes in rodent olfactory preferences, converting an innate aversion for cat odor into attraction, in order to enhance their own transmission^[15]^. Nevertheless, Boillat *et al* found that *T. gondii* infection could commonly shift the host’s aversion to predators, which are not specific to cats. And this alternation may be related to cysts in the host brain^[16]^. As far as humans are concerned, a large amount of seroepidemiological data have shown that *T. gondii* infection has increased the incidence of mental illness, especially focus on the relationship among schizophrenia, suicide and traffic accidents^[17-20]^.

*T. gondii* can infect a variety of cells in the CNS, including neuronal cells, astrocytes, microglia and Purkinje cells. Studies have reported that tachyzoites in CNS are more susceptible to astrocytes^[21]^. Astrocytes are the most abundant glial cell type in CNS, constitute a heterogeneous cell population and could maintain neural homeostasis^[22]^. Astrocytes play an important role including regulation of energy metabolism, brain barrier, synaptic structure and plasticity^[23-26]^. They can also interact with CD^4+^ and CD^8+^ T cells to prevent neuronal damage^[27]^. However, regulation details of astrocytes infected with tachyzoites or/and bradyzoites are still unknown. Therefore, exploring the response of astrocytes infected with different stages of *T. gondii* will help to understand the mechanism of interaction between *T. gondii* and host CNS, and lay a solid foundation for the study of acute and chronic Toxoplasmosis.

Therefore in this study, we have established an in vitro infected astrocytes model using *T. gondii RH* strain and mouse C8-D1A astrocytes. Based on the acute infection model with *RH* tachyzoites, we have also successfully transformed the tachyzoites into bradyzoites in C8-D1A cells. By using the Label-free proteome detection, we have figured out the changes of host protein expression before and after infection with tachyzoite and bradyzoite, respectively. Our results revealed the molecular mechanism of astrocytes to parasites at different stages of *T. gondii* infection. It can also provide new insights into the pathogenic mechanism of acute and chronic toxoplasmosis, and show new targets for the development of anti-*T. gondii* therapeutic drugs.

## 2. Materials and methods

### 2.1 *T. gondii* and cell culture

*T. gondii* tachyzoites (*RH* strain) were maintained using human foreskin fibroblast (HFF) cells, which were cultured in DMEM supplemented with 10% fetal bovine serum (FBS) and 100 IU/mL penicillin, and 100 μg/mL streptomycin at 37°C in a humidified atmosphere of 5% CO_2_. When the infected HFF monolayer was lysed, collected cell mixture and filtered through a 5 μm filter to obtain *T. gondii* tachyzoites, counted and stored at -80°C. C8-D1A mouse astrocytes were cultured under the same conditions as HFF cells.

### 2.2 Sample preparation and collection

C8-D1A (5×10^5^ cells) were seeded on 8 mm coverslips in 6-well plates, and cultured in 37°C with 5% CO_2_.When cells formed confluent monolayers, replace medium with 3% FBS. *RH* strain of *T. gondii* were added to C8-D1A (The ratio of cells to tachyzoites was 10:1) for 24 h. PBS washed two times and collected samples, centrifuged at 1000 g for 5 minutes, stored at -80°C, the group was named C8-TA. Alkaline-induced transformation of tachyzoites in vitro were performed. The *T. gondii* tachyzoites were added to cells at a ratio of 1:10 and cultured for 8 h at 37°C with 5% CO_2_, then continued to culture for 96 h in alkaline media (pH 8.2) and changed the medium every 1-2 days. PBS washed two times and collected samples, centrifuged at 1000 g for 5 minutes, stored at -80°C, the group was named C8-BR. The uninfected group was named C8. There were three replicates for each group, therefore 9 samples were used for following experiment.

### 2.3 RT-PCR

RT-PCR was used to detect the expression of bradyzoite antigen 1 (BAG1). Total RNA was extracted from C8-BR group with TRIzol (Sigma-Aldrich, USA). The isolated RNA was converted to cDNA using Takara PrimerScriptTM RT reagent Kit according to the instructions. BAG1 primers were as follows:

Forward primer: 5’-TCGCCTCTCAACAGCTAGAC-3’;

Reverse primer: 5’-CCCTGAATCCTCGACCTTGAT-3’;

The reaction conditions were 94 °C for 5 min, 94 °C for 40 s, 56 °C for 40 s, and 72 °C for 1 min. PCR products were analyzed by 1% agarose gel electrophoresis.

### 2.4 Protein extraction and trypsin digestion

All of the samples were sonicated three times on ice using a high intensity ultrasonic processor (Scientz) in lysis buffer (8 M urea, 1% Protease Inhibitor Cocktail). Centrifuged at 12,000g at 4°C for 10 min to remove remaining debris. BCA kit was used to determine the protein concentration according to the manufacturer’s instructions. For digestion, the protein solution of each sample was reduced with 5 mM dithiothreitol for 30 min at 56 °C and then alkylated with 11 mM iodoacetamide in the dark for 15 min at room temperature. 100 mM TEAB was added to dilute the sample. Finally, trypsin was added for the first digestion overnight (trypsin-to-protein mass ratio was 1:50) and performed a subsequent 4 h-digestion (trypsin-to-protein mass ratio was 1:100).

### 2.5 LC-MS/MS Analysis

The tryptic peptides were dissolved in 0.1% formic acid (solvent A), directly loaded onto a home-made reversed-phase analytical column (15-cm length, 75 μm i.d.). The gradient was comprised of an increase from 6% to 23% solvent B (0.1% formic acid in 98% acetonitrile) over 26 min, 23% to 35% in 8 min and climbing to 80% in 3 min then holding at 80% for the last 3 min, all at a constant flow rate of 400 nL/min on an EASY-nLC 1000 UPLC system. The peptides were subjected to NSI source followed by tandem mass spectrometry (MS/MS) in Q ExactiveTM Plus (Thermo) coupled online to the UPLC. The electrospray voltage applied was 2.0 kV. The m/z scan range was 350 to 1800 for full scan, and intact peptides were detected in the Orbitrap at a resolution of 70,000. Peptides were then selected for MS/MS using NCE setting as 28 and the fragments were detected in the Orbitrap at a resolution of 17,500. A data-dependent procedure that alternated between one MS scan followed by 20 MS/MS scans with 15.0s dynamic exclusion. Automatic gain control (AGC) was set at 5E4.

### 2.6 Database Search

The resulting MS/MS data were processed using Maxquant search engine (v.1.5.2.8). Tandem mass spectra were searched against Mus musculus data in the Uniprot database concatenated with reverse decoy database. Trypsin/P was specified as cleavage enzyme allowing up to 2 missing cleavages. Mass tolerance for precursor ions was set as 20 ppm in First search and 5 ppm in Main search, and the mass tolerance for fragment ions was set as 0.02 Da. Carbamidomethyl on Cys was specified as fixed modification, oxidation on Met was specified as variable modifications. Label-free quantification method was LFQ, FDR was adjusted to < 1% and the minimum score for peptides was set > 40.

### 2.7 Bioinformatic analysis

Wolfpsort (http://www.genscript.com/psort/wolf_psort.html) was used to predicate subcellular localization of DEPs. Eukaryotic orthologous group (http://genome.jgi.doe.gov/help/kogbrowser.jsf) was performed on all DEPs for further functional classification by aligning their sequences with the KOG database. The UniProt-GOA database (http://www.ebi.ac.uk/GOA/) together with InterProScan soft (http://www.ebi.ac.uk/InterProScan/) were used to analyze biological process, cellular component and molecular function of DEPs. The Kyoto Encyclopedia of Genes and Genomes (KEGG) database(https://www.genome.jp/kegg) was used to analyze signaling pathways involved in DEPs. A two-tailed Fisher’s exact test was employed to test the enrichment of the differentially expressed protein against all identified proteins. A corrected *p*-value < 0.05 was considered significant in database analysis. Cluster membership was visualized by a heat map using the “heatmap.2” function from the “gplots” R-package.

### 2.8 Parallel Reaction Monitoring (PRM) Validation

To verify the accuracy of Label-free proteome quantification analysis, we selected 18 DEPs proteins for PRM assay. The methods of protein extraction and trypsin digestion were as described above. In LC-MS/MS Analysis, the gradient was comprised of an increase from 6% to 23% solvent B (0.1% formic acid in 98% acetonitrile) over 38 min, 23% to 35% in 14 min and climbing to 80% in 4 min then holding at 80% for the last 4 min, all at a constant flow rate of 700 nL/min on an EASY-nLC 1000 UPLC system. Automatic gain control (AGC) was set at 3E6 for full MS and 1E5 for MS/MS. The maxumum IT was set at 20 ms for full MS and auto for MS/MS. The isolation window for MS/MS was set at 2.0 m/z. Peptide parameters were as follows: enzyme was set as Trypsin [KR/P], Max missed cleavage set as 2. The peptide length was set as 8-25. precursor charges were set as 2, 3, ion charges were set as 1, 2, ion types were set as b, y, p. The product ions were set as from ion 3 to last ion, the ion match tolerance was set as 0.02 Da.

## 3. Results

### 3.1 Identification of DEPs between C8-TA, C8-BR and C8 groups

We used Label-free proteome to quantitatively analyze the host proteins and in tachyzoite and tachyzoite to bradyzoite transformation stages of *T. gondii* infection (Supplementary Figure 1). Among the 9 samples, a total of 32906 peptides and 30808 unique peptides were identified. We identified 4676 proteins as host protein, of which 3415 proteins were quantified (Supplementary Table S1). We defined fold change≥1.5 or ≤0.67 and *p*-value≤0.05 as the criteria to analyze the DEPs of C8-TA, C8-BR and C8 group (Figure 1A). There were a total of 163 DEPs, of which 109 were up-regulated and 54 were down-regulated in C8-TA group compared with C8 group (Supplementary Table S2A, Figure 1B). In addition, there were 719 DEPs, 495 host proteins were up-regulated and 224 host proteins were down-regulated in C8-BR group compared with C8-TA group (Supplementary Table S2B, Figure 1C).

**Figure 1.**
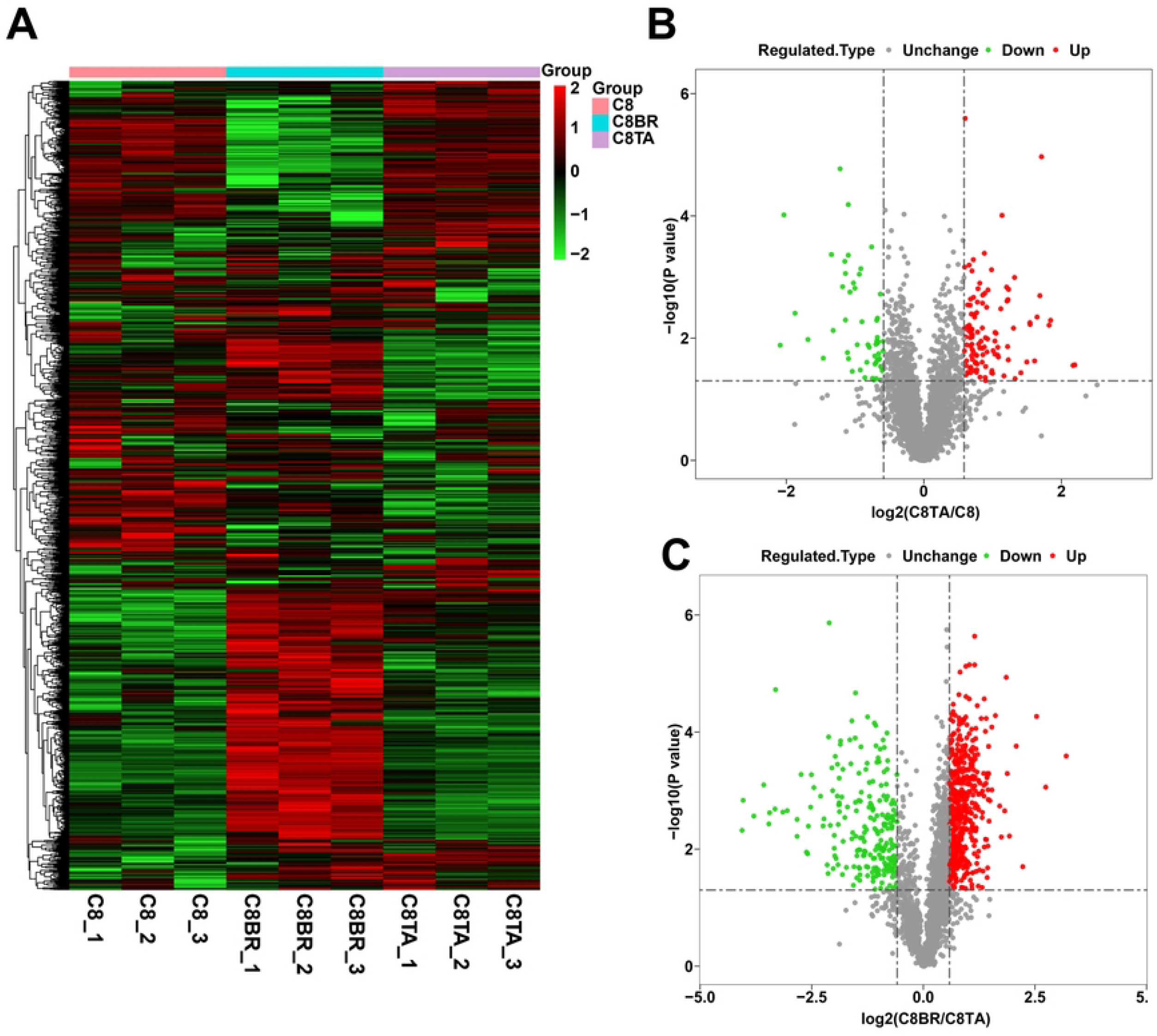
Proteome analysis of DEPs in tachyzoite and tachyzoite to bradyzoite transformation stages of *T. gondii* infection. (A) Heat map shown the DEPs in the three groups. (B) Volcano plot shown DEPs between the C8-TA group and the C8 group. (C) Volcano plot shown DEPs between the C8-BR group and the C8 group.

### 3.2 Tachyzoite infection altered multiple immunoregulatory processes in astrocytes

To investigate the biological functions of the DEPs between the C8-TA group and the C8 group, we firstly analyzed the subcellular localization of the DEPs. As shown in Figure 2A, in the C8-TA vs C8 group, 38.27% DEPs were located in cytoplasma, 14.81% DEPs were located in plasma membrane, and 17.28% DEPs were located in nucleus. KOG (EuKaryotic orthologous groups) was used to predict the potential functions of DEPs in the C8-TA vs C8 group (Supplementary Table S3, Figure 2B). The results showed that the top five KOG classifications were [U] Intracellular trafficking, secretion, and vesicular transport, [J] Translation, ribosomal structure and biogenesis, [O] Posttranslational modification, protein turnover, chaperones, [Z] Cytoskeleton, [A] RNA processing and modification. GO enrichment analysis was performed for the functional annotation of DEPs including three categories: biological process (GO-BP), cellular compartment (GO-CC) and molecular function (GO-MF) (Supplementary Table S4A, Figure 2C). We focused on the biological processes and found that the DEPs mainly enriched in immune-related biological processes, such as defense response to other organism, defense response to virus, response to interferon-beta, innate immune response, detection of virus, which indicated that the DEPs mainly involved in immune regulations.

**Figure 2.**
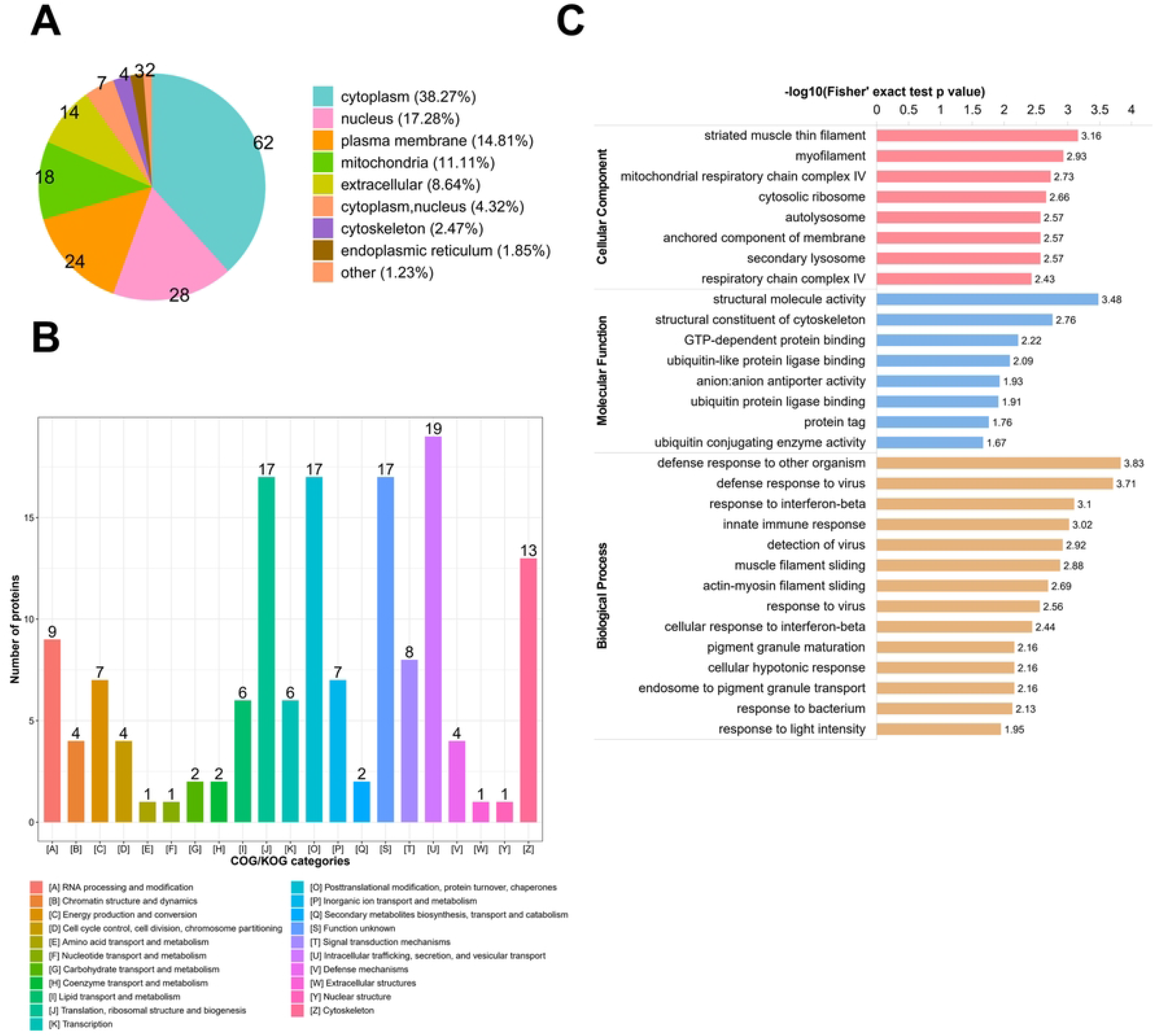
Location and functional classification of DEPs between the C8-TA group and the C8 group. (A) Location of subcellular structures of DEPs between the C8-TA group and the C8 group. (B) KOG classification of DEPs between the C8-TA group and the C8 group. (C) GO enrichment analysis of DEPs between C8-TA vs C8 group included biological process, cellular composition, and biological function.

Next, we comprehensively analyzed the biological functions of up-regulated and down-regulated DEPs in C8-TA vs C8 group. The GO-BP results showed that in C8-TA vs C8 group, the up-regulated DEPs were mainly involved in the following biological processes: cellular hypotonic response, response to light intensity, learning, ribosomal small subunit biogenesis, negative regulation of transporter activity (Supplementary Table S4B, Figure 3A). The down-regulated DEPs were mainly related to immune responses, such as defense response to other organisms, defense response to virus, response to interferon-beta, response to virus, and innate immune response (Supplementary Table S4C, Figure 3B). KEGG enrichment analysis of up-regulated and down-regulated DEPs showed the signaling pathways during the infection of *T. gondii* tachyzoites (Supplementary Table S5A, 5B). As shown in Figure 3C, D, the up-regulated proteins were mapped to 18 signaling pathways, and the down-regulated proteins were mapped to 10 signaling pathways in C8-TA vs C8 group. As expected the down-regulated proteins were enriched in inflammatory signaling pathways, such as Herpes simplex virus 1 infection, RIG-I-like receptor signaling pathway, Amoebiasis, Systemic lupus erythematosus. The above results indicated that the host immune response was activated during the invasion of *T. gondii*, interestingly, our results showed that the down-regulated proteins could play a major role in immune regulation, which was different from the biological traits of *T. gondii* invaded epithelial cells, indicated that when tachyzoites invaded astrocytes for 24 h, the immune response of the host was gradually weakened. This might be a protective measurement for *T. gondii* to avoid host astrocytes generating strong neuro-immune response.

**Figure 3.**
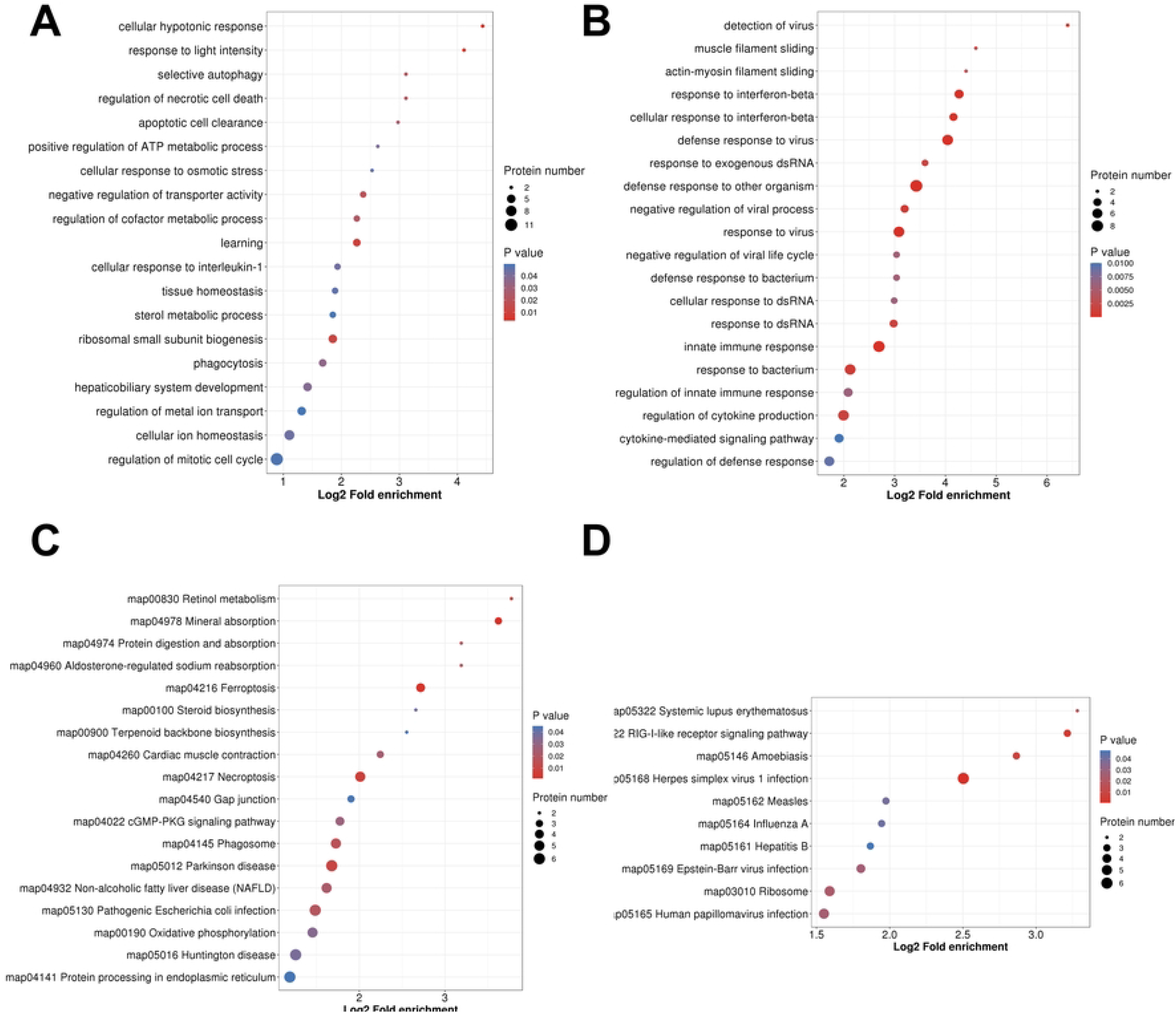
GO and KEGG pathway enrichment analysis of up-regulated and down-regulated DEPs between C8-TA group and C8 groups. (A) GO enrichment analysis of up-regulated DEPs. (B) GO enrichment analysis of down-regulated DEPs. (C) KEGG pathway enrichment analysis of up-regulated DEPs. (D) KEGG pathway enrichment analysis of down-regulated DEPs.

### 3.3 The conversion of the tachyzoites into bradyzoites upregulated glucose metabolism of astrocytes

Tachyzoite to bradyzoite differentiation is a key aspect of *T. gondii* biology and pathogenesis. To better understand the mechanism of interaction between *T. gondii* and the host during differentiation of *T. gondii*, we performed a comprehensive bioinformatics analysis in C8-BR vs C8-TA group. First, we analyzed the sublocalization of all DEPs by WoLF PSORT software (Figure 4A). The results showed that 33% of the DEPs were localized in cytoplasm, 26% in nucleus, and 17% in mitochondria. In up-regulated DEPs, most of them were located in the cytoplasm (36%), followed by nucleus (21.41%) and mitochondria (20.61%) (Figure 4B). In down-regulated DEPs, most of them were located in the nucleus (37%), followed by cytoplasm (26%) and plasma membrane (11%) (Figure 4C).

**Figure 4.**
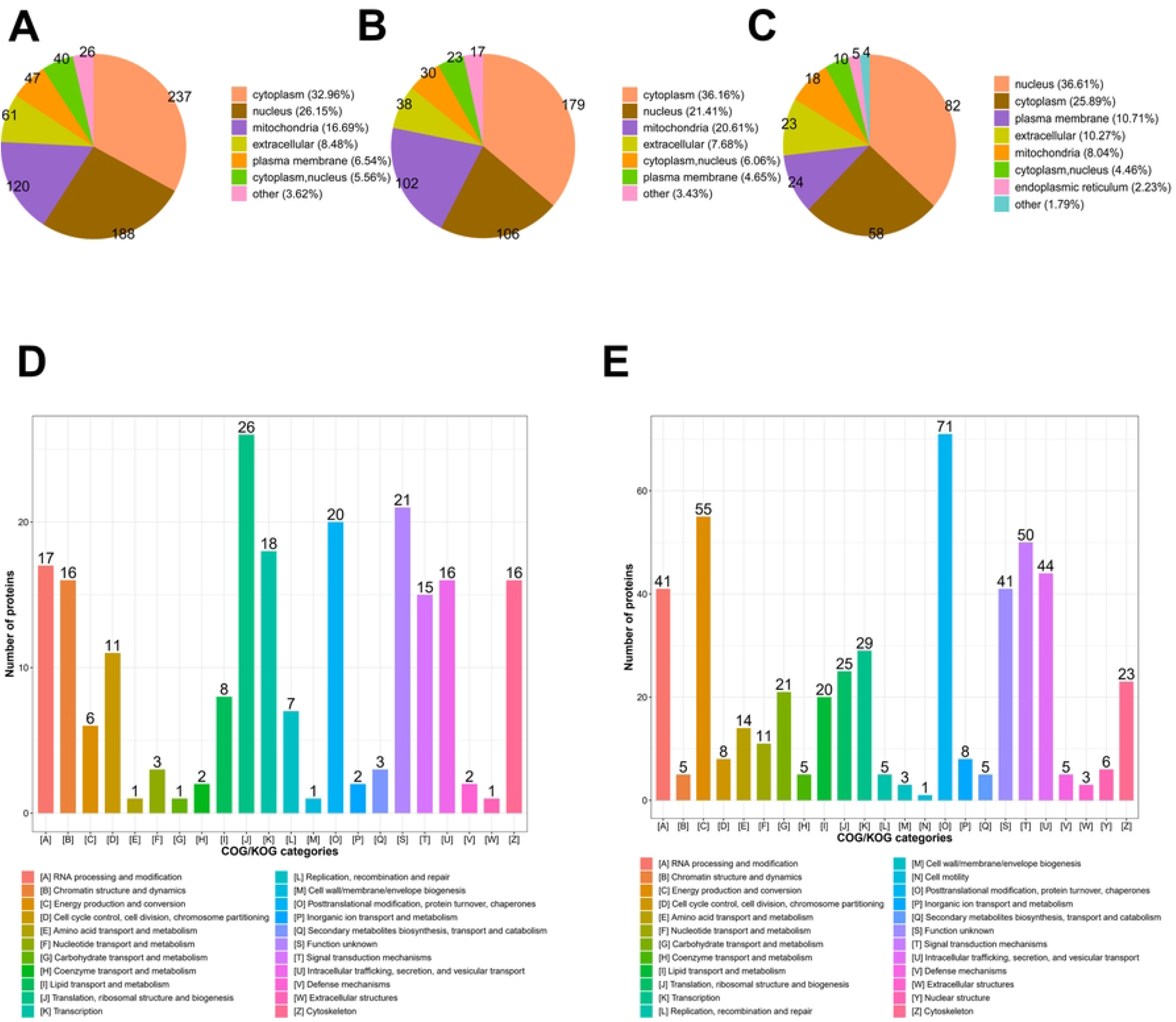
Subcellular localization and functional classification of DEPs between C8-BR group and C8-TA groups. (A) The location of subcellular structures of all DEPs. (B) The location of subcellular structures of up-regulated DEPs. (C) The location of subcellular structures of down-regulated DEPs. (D) The KOG classification of up-regulated DEPs. (E) The KOG classification of down-regulated DEPs.

We used KOG database to predict the function and classification of up and down-regulated proteins (Supplementary Table S6A, 6B, Figure 4D, E). The results showed that the functions of up-regulated DEPs were consistent with down-regulated DEPs, and the DEPs were mainly enriched in the following categories: [O] Posttranslational modification, protein turnover, chaperones, [T] Signal transduction mechanisms, [C] Energy production and conversion, [U] Intracellular trafficking, secretion, and vesicular transport, [A] RNA processing and modification pathway, suggested that during the chronic infection stage, epigenetic mechanisms may mediate the effects on host nutrient metabolism in the coexistence of host and bradyzoites. GO enrichment analysis showed that the DEPs were mainly involved in metabolism-related biological processes including dicarboxylic acid metabolic process, oxidoreduction coenzyme metabolic process, hexose biosynthetic process, monocarboxylic acid metabolic process, tricarboxylic acid metabolic process (Supplementary Table S7A, Supplementary Figure 2A). Among them, glucose metabolism related biological processes were up-regulated, such as dicarboxylic acid metabolic process, oxidoreduction coenzyme metabolic process, carboxylic acid metabolic process, tricarboxylic acid metabolic process, nucleoside phosphate metabolic process (Supplementary Table S7B, Figure 5A). However, there were no significant biological processes enrichment for down-regulated proteins, the top five categories of down-regulated proteins were protein localization to endoplasmic reticulum, ribosomal large subunit biogenesis, peptide biosynthetic process, DNA conformation change, peptide metabolic process (Supplementary Table S7C, Figure 5B).

**Figure 5.**
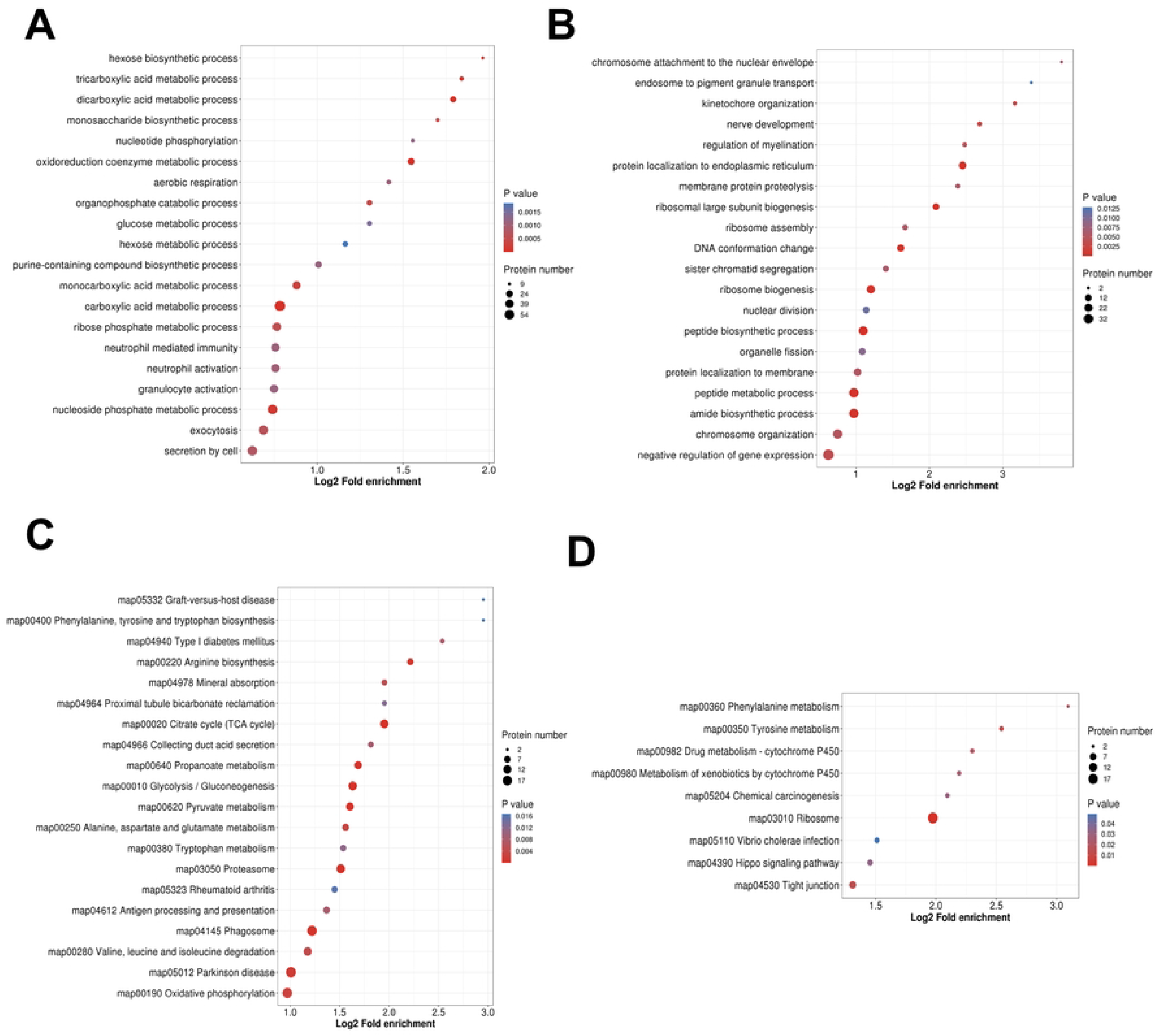
GO and KEGG pathway enrichment analysis of up-regulated and down-regulated DEPs between C8-BR group and C8-RH groups. (A) GO enrichment analysis of up-regulated DEPs. (B) GO enrichment analysis of down-regulated DEPs. (C) KEGG pathway enrichment analysis of up-regulated DEPs. (D) KEGG pathway enrichment analysis of down-regulated DEPs.

KEGG pathway enrichment was used to identify pathways of DEPs (Supplementary Table S8A, Supplementary Figure 2B), the results showed that DEPs significant enriched in metabolism-related pathways, the top five were Glycolysis/Gluconeogenesis, Citrate cycle (TCA cycle), Phenylalanine metabolism, Phagosome, Pyruvate metabolism. We further investigated the function of up and down-regulated DEPs and found that up-regulated proteins were mainly involved in metabolism-related signaling pathways including Citrate cycle (TCA cycle), Glycolysis/Gluconeogenesis, Proteasome, Phagosome, Pyruvate metabolism, which was consistent with the GO-BP (Supplementary Table S8B Figure 5C). The part of down-regulated proteins involved in amino acid metabolism, such as Tyrosine metabolism, Phenylalanine metabolism (Supplementary Table S8C, Figure 5D). During the transformation from tachyzoite infection stage to bradyzoite infection stage, the function of host proteins may have undergone a gradual transition from immune emergency mode to up-regulation metabolic pathways. The effects on host metabolism during transformation suggested that, the increased host’s metabolism could accelerate the decomposition of glucose, so the metabolism of *T. gondii* was decreased, which was more conducive to its long-lived in their hosts.

### 3.4 Verification of differentially expressed proteins by PRM

To evaluate the accuracy of Label-free proteome quantification techniques, a total of 18 DEPs were selected for PRM analysis. Based on GO and KEGG annotations, we chose Bystin (Bysl), Fibronectin (Fn1), Signal transducer and activator of transcription 1 (Stat1), Serpin B6 (Serpinb6), Guanylate-binding protein 4 (Gbp4), Interferon-induced protein with tetratricopeptide repeats 3 (Ifit3), Antiviral innate immune response receptor RIG-I (Ddx58), 60S ribosomal protein L23 (Rpl23), 40S ribosomal protein S24 (Rps24) in C8-TA vs C8-group and Dihydrolipoyl dehydrogenase (Dld), Aspartate aminotransferase, cytoplasmic (Got1), DNA (cytosine-5)-methyltransferase 1 (Dnmt1), Nischarin (Nisch), Enoyl-CoA hydratase, mitochondrial (Echs1), Dihydrolipoyllysine-residue acetyltransferase component of pyruvate dehydrogenase complex (Dlat), Tricarboxylate transport protein (Slc25a1), NAD-dependent protein deacetylase sirtuin-2 (Sirt2), Omega-amidase NIT2 (Nit2) in C8-BR vs C8-TA group (Figure 6, Supplementary Table S9). The results showed that the changing trends of 18 DEPs protein in PRM were consistent with Label-free proteome quantification, suggesting that Label-free proteome quantification outcomes were relatively reproducible and reliable.

**Figure 6.**
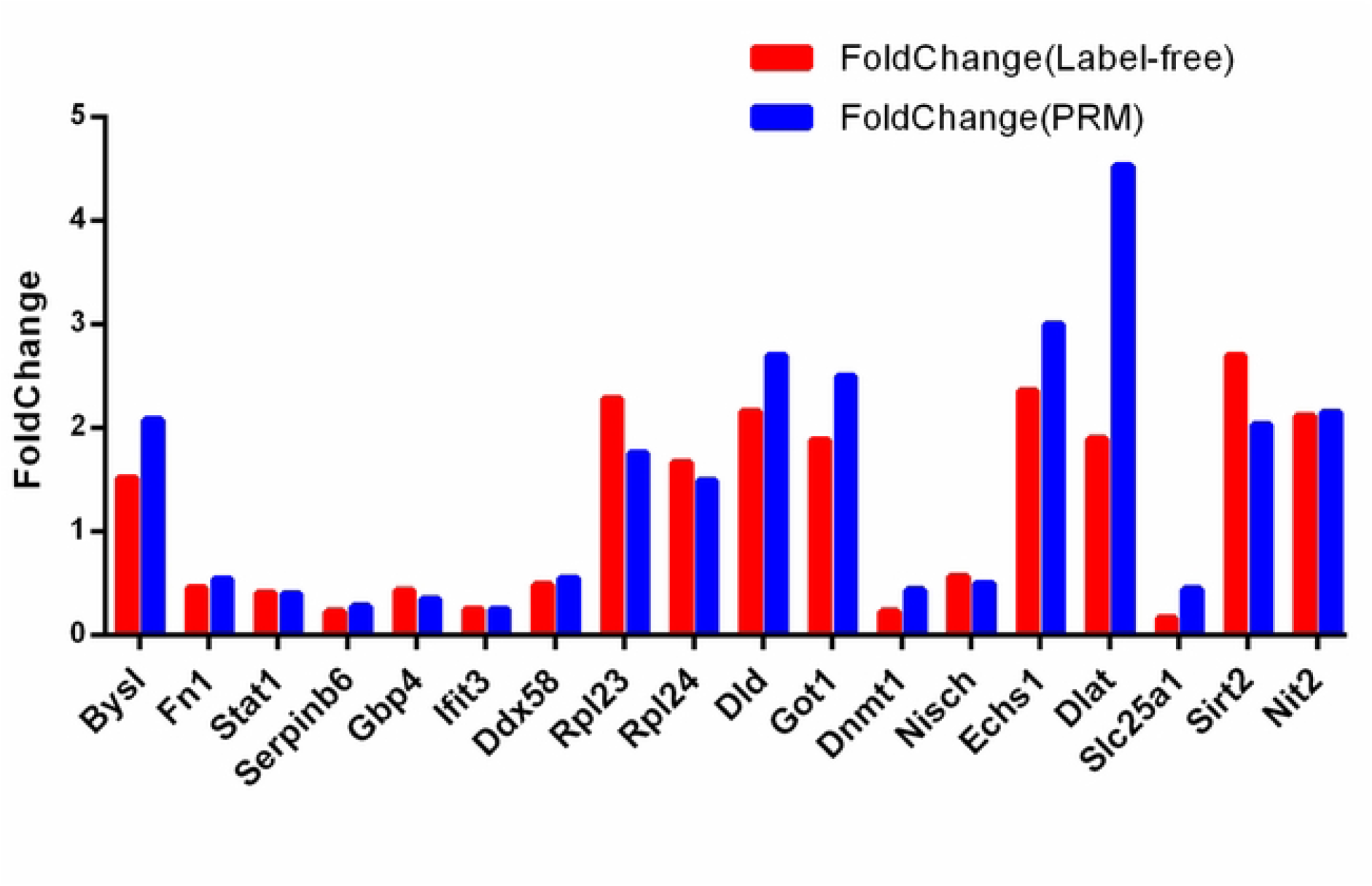
Comparative analyses of label-free proteomics and PRM results for 18 screened DEPs.

## 4. Discussion

The invasion of *T. gondii* tachyzoites and bradyzoites in the host causes acute infection and chronic infection respectively, resulting in clinical manifestations of toxoplasma encephalopathy, chorioretinitis, miscarriage, stillbirth and schizophrenia^[17,28-30]^. Since there is no effective method to eliminate tissue cysts up to now, the damage of host CNS caused by *T. gondii* accompany with mental and behavioral disorders has been brought into focus. Therefore, understanding the proteomics changes in host neurogliocytes after tachyzoites/bradyzoites infection would identify the negative effects of different stages on the host, and provide targets for developing new vaccines and drugs to against brain damages caused by *T. gondii*.

In the present study, our results showed that the DEPs in the acute infection stage were mainly enriched in immune-related biological processes, while the DEPs in the chronic infection stage were mainly enriched in metabolic-related biological processes. Therefore our results suggested that *T. gondii* would regulate host neurogliocytes by distinct mode in the two stages. Similarly, previous researches on tachyzoite infection stage indicated that the up-regulated DEPs were also involved in immune inflammation related pathways to prevent acute infection^[31]^, while those similar studies mainly based on epithelial cells such as HFF cells.

As we know, invasion of *T. gondii* can cause inflammation in the host, and astrocytes have been proved to be a pivotal regulator of CNS inflammatory responses^[32]^. The borders and scars of astrocyte could serve as functional barriers that restrict the entry of inflammatory cells into CNS parenchyma in health and disease, simultaneously, it also has powerful pro-inflammatory potential. Therefore, we have established acute/chronic infection models with mouse astrocytes. Interestedly, although our DEPs results in tachyzoites infection models were also involved in immune inflammation related pathways, changes in protein expression were very different to the results of previous studies in epithelial cells.

In our study, the down-regulation DEPs rather than up-regulation DEPs were involved in immune regulation-related processes, especially in defense response pathways and innate immune response pathways. Consistently, Cekanaviciute reported that after *T. gondii* infection in host CNS, the immune function must be restricted to prevent excessive neuronal damage, in order to keep a balance among *T. gondii*, brain and the immune system^[33]^. As a result, we have focused on four down-regulated and immune-related DEPs including Stat1, Gbp4, Ifit3 and Ddx58 after tachyzoite infection. Stat1 is an important immune inflammatory factor in CNS and is involved in the immune regulation of various cells. Studies have shown that inhibition of Stat1 can promote bradyzoite formation^[34]^. In this study, the expression of Stat1 was significantly decreased, suggesting that after tachyzoite infection in astrocytes, the down-regulated Stat1 might participate in promoting the transformation of tachyzoites to bradyzoites, as well as inhibiting the persistent infection of tachyzoites. Gbp4 and Ifit3 are IFN-γ-inducing proteins. Hu *et al* found that Gbp4 could negatively regulate virus-induced type I IFN and antiviral responses by interacting with IFN regulatory factor (IRF) 7 during viral infection, thus the following researches on their roles in astrocytes during *T. gondii* infection may reveal a new mechanism by which astrocytes against parasites^[35]^. Ddx58 is also known as RIG-I, can function as an innate antiviral immune response receptor and play an important role in antiviral innate immunity^[36-38]^. In vesicular stomatitis virus (VSV)-infected astrocytes, RIG-I knockdown significantly reduced inflammatory cytokine production in astrocytes^[39]^, and so the down regulation of Ddx58 may be also involved in the transformation of tachyzoites to bradyzoites in host CNS.

The conversation from tachyzoites to bradyzoites of *T. gondii* is a key to establish chronic infection and an important link in the pathogenesis of *T. gondii*. We comprehensively analyzed the DEPs between the bradyzoites infection group and the tachyzoite infection group. The results showed that the DEPs were mainly involved in metabolism-related biological processes and signaling pathways, and the detailed GO and KEGG analyses indicated that the host’s glucose metabolism and a part of amino acid metabolism process have been changed significantly during chronic infection. Further, we found that the up-regulated DEPs were mainly involved in glucose metabolism and entered the TCA process, especially for Echs1.

Echs1 is a key enzyme involved in the metabolism of fatty acyl-CoA esters^[40]^. In fatty acid β-oxidation, it could increase the synthesis of acetyl-CoA and promoted the TCA cycle, thereby increasing the process of glucose metabolism. It has been reported that Echs1 deficiency (Echs1D) leaded to the impaired ATP production and metabolic acidosis in patients^[41]^. As an obligate intracellular parasite, *T. gondii* obtains all nutrients from host cells to support its intracellular growth and proliferation. Glucose and glutamine are raw materials for tachyzoites to complete the classic TCA and then synthesized ATP^[42,43]^. Therefore the increase of host glucose metabolism would accelerate the decomposition of glucose, result in the decreasing uptake of glucose by *T. gondii*. In the present study, the up-regulated Echs1 may be an important host receptor of invasion parasites, and associate with the inhibition of parasites self-glucose metabolism, thereby promote the formation of intracellular bradyzoites and establish long-term latent infection in the host.

Neuronal degeneration caused by chronic infection of *T. gondii* is an important pathogenesis of neurodegenerative diseases, but the mechanism has not been fully elucidated. Our results also found important clues and potential targets for this process, such as Dld, Sirt2 and Got1.

Dld, also known as Dihydrolipoyl dehydrogenase, is a mitochondrial enzyme that is essential for eukaryotic cell metabolism^[44]^. Ahmad reported that Dld was related to Alzheimer’s disease(AD), and inhibition of Dld expression would lead to a significant recovery of Aβ pathological degradation. This could be explained that the inhibition of Dld would down-regulate metabolism-related signaling pathways and reduce the host’s energy metabolism, which would be beneficial to alleviate the symptoms of AD^[45]^. In this study, we found that Dld was significantly up-regulated in bradyzoites infection group, which may provide a new insight for explaining the mechanism of *T. gondii* infection-induced Alzheimer’s disease. In addition, Dld is also associated with severe diseases in infants, causing developmental delay, hypotonia and metabolic acidosis^[46]^. Recent studies have shown that Leishmania’s self-encoded Dld induced a protective cellular immune response in L. major-infected mice, which could serve as a design site for Duchenne vaccines for kala-azar prevention^[24,47]^.

Sirt2 is an NAD+-dependent deacetylase that is widely involved in cell division, angiogenesis^[48-50]^, energy metabolism^[51]^ and neurodegenerative^[52,53]^, cardiovascular disease^[54,55]^, oxidative stress^[56]^and many cancers^[57,58]^, etc. Studies have found that Sirt2 was the most abundant sirtuin expressed in mammalian CNS, especially in cortex, striatum, hippocampus, and spinal cord, suggesting that it might have a role in CNS^[59]^. Dopaminergic (DA) neurons plays a vital role in CNS, and DA neuron hyperfunction was involved in several neurological disorders including schizophrenia and Parkinson’s Disease. Sirt2 expression was dramatically increased during the differentiation of human embryonic stem cells (hESCs) into midbrain DA neurons^[60]^. In addition, Sirt2 knockout (Sirt2−/−) mice displayed fewer DA neurons and less dense striatal fibers in the substantia nigra^[61]^. Sirt2 inhibition also improves cognitive impairment in different AD animal models and promotes neuronal survival^[62,63]^. This suggested that Sirt2 can be a potential therapeutic target for neurodegenerative disease. Moreover, Sirt2 is also involved in ATP synthesis. Recent studies have shown that Sirt2 localized to the inner mitochondrial membrane of the mouse brain, and mice lacking Sirt2 showed decreased ATP levels in the striatum^[64]^. In our study, the expression of Sirt2 was up-regulated after chronic infection, which is consistent with Mcconkey’s report, they found that dopaminergic cells and brain tissues encysted with cerebral cysts have increased levels of dopamine synthesis and release^[65]^. Moreover, treatment of rats and mice with dopamine receptor antagonists could inhibit the behavioral changes induced by *T. gondii* infection^[66,67]^. In addition, some parasites such as Leishmania, T. brucei, and Schistosoma could encode Sirt2 homologous protein, and more importantly, it was essential for parasite growth^[68-70]^. Thus Sirt2 might serve as a potential treatment target for psychiatric disorders induced by manipulative parasites.

Neuroendocrine programs and neurotransmitter imbalance may act as the physiological basis for *T. gondii* induced psychiatric and behavioral disorders^[71]^. Glutamate (GLU) is the most abundant neurotransmitter in the brain, and its excitability plays a crucial role in brain structure and function. Got1 aspartate aminotransferase is a type of aminotransferase that catalyzes the transamination of aspartate and α-ketoglutarate to form glutamate and oxaloacetate. Glutamate dehydrogenase (Glud1) is a mitochondrial enzyme that catalyzes the reductive fixation of ammonia to α-ketoglutarate to form glutamate. Both Got1 and Glud1 expression were found to be up-regulated in our study, suggesting that the glutamate synthesis might be increased in the chronic infection group. Our finding is also consistent with a recent study, which showed that chronic infection with *T. gondii* could cause an increase in extracellular glutamate and a two-fold decrease in glutamate transporter expression in glial cells^[72]^. As hyperexcitability of GLU is neurotoxic and leading to brain damage, neurological disorders (eg, ALS, multiple sclerosis, AD, Huntington’s disease, Parkinson’s disease) and psychiatric disorders (eg, schizophrenia, depression, bipolar disorder). Our finding provides new evidence to explain the host mental behavioral disorders induced by chronic *T. gondii* infection.

In conclusion, our results systematically analyzed the proteomic changes in astrocytes infected with not only tachyzoites, but also bradyzoites, respectively. We surprisingly discovered that *T. gondii* tachyzoites can cause down-regulation of immune-related pathways in astrocytes, and the inhibited expression of Echs1 might be associated with the transformation of tachyzoites to bradyzoites. However, during the bradyzoites infection stage, metabolism rather than immune pathways of the host was changed significantly. Both the glucose metabolism pathways and the expression of some metabolism-related enzymes in astrocytes were significantly up-regulated, such as Dld, Sirt2 and Got1. Since their expression was closely related to chronic degenerative diseases and psychiatric diseases, which could provide a new explanation for host mental and behavioral disorders induced by chronic infection of *T. gondii*.

## Author Contributions

HX and KY. conceived the project and designed the experiments. HX, CX, GZ, LZ, HX and LD performed the experiments. HX, KY, CX analyzed the data. HS, QW, JZ and JH helped to design the layout of the pictures. HX wrote the manuscript and KY revised the manuscript. All authors contributed to the article and approved the submitted version.

## Figure legends

**Supplementary Figure 1 Agarose gel electrophoresis of PCR product for BAG1 gene**.

**Supplementary Figure 2 GO and KEGG enrichment analysis of all DEPs between C8-BR group and C8-RH group**. (A) GO enrichment analysis of all DEPs. (B) KEGG enrichment analysis of all DEP.

